# PAVR: High-Resolution Cellular Imaging via a Physics-Aware Volumetric Reconstruction Framework

**DOI:** 10.64898/2026.03.04.709609

**Authors:** Xuanwen Hua, Keyi Han, Zhi Ling, Olivia Reid, Zijun Gao, Hongmanlin Zhang, Edward A. Botchwey, Parvin Forghani, Wenhao Liu, Mithila Anil Sawant, Afsane Radmand, Hyejin Kim, James E Dahlman, Aparna Kesarwala, Chunhui Xu, Shu Jia

**Affiliations:** Laboratory for Systems Biophotonics, Georgia Institute of Technology, Atlanta, Georgia, USA; Wallace H. Coulter Department of Biomedical Engineering, Georgia Institute of Technology and Emory University, Atlanta, Georgia, USA; Parker H. Petit Institute for Bioengineering and Biosciences, Georgia Institute of Technology, Atlanta, Georgia, USA; George W. Woodruff School of Mechanical Engineering, Georgia Institute of Technology, Atlanta, Georgia, USA; Department of Pediatrics, School of Medicine, Emory University, Atlanta, Georgia, USA; School of Materials Science and Engineering, Georgia Institute of Technology, Atlanta, Georgia, USA; Department of Radiation Oncology, School of Medicine, Emory University, Atlanta, Georgia, USA; Winship Cancer Institute, School of Medicine, Emory University, Atlanta, Georgia, USA; Department of Chemical Engineering, Georgia Institute of Technology, Atlanta, Georgia, USA

## Abstract

The rapid convergence of advanced microscopy and deep learning is transforming cell biology by enabling imaging systems in which optical encoding and computational inference are jointly optimized for volumetric information capture and interpretation. However, broadly accessible three-dimensional imaging at high spatiotemporal resolution remains constrained by volumetric reconstruction throughput, susceptibility to artifacts, and the burden of collecting modality-matched training data. Here, we introduce PAVR, a physics-aware light-field imaging platform that integrates single-shot volumetric acquisition with fast, end-to-end volumetric reconstruction. PAVR is trained entirely using *in silico* system responses, avoiding reliance on external high-resolution ground-truth modalities and enabling sample-independent reconstruction across diverse biological contexts. Using fixed and live mammalian cells, we demonstrate multicolor volumetric imaging of subcellular organelles, three-dimensional tracking of autofluorescent particles, and high-speed visualization of organelle remodeling and interactions. We further extend PAVR to quantify coupled morphological and functional dynamics in beating human induced pluripotent stem cell–derived cardiomyocytes under pharmacological perturbation. Together, PAVR establishes a scalable hardware–software platform for high-throughput volumetric imaging and quantitative analysis of dynamic cellular systems in both basic and translational settings.

## INTRODUCTION

Optical microscopy has fundamentally transformed modern cell biology by enabling direct visualization of the molecular and structural foundations of life with high sensitivity, molecular specificity, and spatiotemporal precision^1-3^. As biological questions continue to expand in complexity, scale, and dimensionality, computational approaches have become indispensable complements to experimental image acquisition^4,5^. In particular, the rapid emergence of deep learning has been transformative, establishing data-driven frameworks that integrate imaging physics with machine intelligence^6^. This paradigm now per-meates the entire microscopy workflow, from optical design^7^ and image formation^8^ to visualization^9^, quantitative analysis^10^, and closed-loop automation^11^. Collectively, these approaches have enabled the extraction of subtle biological features, integration of high-dimensional multimodal datasets^12,13^, and scalable interrogation across diverse biological^14-17^ and translational^18-21^ contexts.

Indeed, the evolution of microscopy modalities, in convergence with deep learning, is redefining the attainable limits of resolution, speed, and sensitivity^22-25^. Advances in computational super-resolution^26^, real-time image reconstruction^27^, and photon-efficient low-light imaging^28^ exemplify the rapidly expanding scope of computation-empowered microscopy hardware systems. Within this landscape, light-field techniques, especially recent Fourier light-field microscopy (FLFM), have drawn increasing attention for infrastructure development and cell biology research, owing to their unique imaging capabilities and accessible implementations based on epi-fluorescence settings^29^. In particular, FLFM captures uniform 4D spatio-angular optical fields, facilitating the recovery of volumetric information without the need for sequential scanning^30-33^. This capability enables high-speed volumetric imaging with reduced photo-damage, high fidelity, and high scalability, supporting investigations spanning single molecules^34,35^ and living cells^36-41^ to tissues^32,33,42,43^ and whole organisms^44^.

As an inherently computation-intensive technique, light-field image reconstruction has traditionally relied on physics-based models formulated using either ray optics^30,31,45^ or wave optics principles^32,33,46^. While ray-optics approaches offer computational efficiency, wave-optics formulations provide improved spatial resolution and image quality, albeit at substantially increased computational cost. Consequently, these trade-offs, together with reconstruction artifacts and limited throughput, remain major constraints for high-resolution, real-time study of dynamic cell biological systems. Contrary to model-based methods, deep learning offers a compelling alternative by exploiting nonlinear mappings between measurements and reconstructions, enabling accelerated image formation with enhanced reconstruction quality^47^. Existing deep learning-based efforts have adopted either fully data-driven, end-to-end formulations for direct 3D inference from raw light-field images^48,49^, or physics-guided, modular neural network components embedded within processing pipelines^50^. Despite these advances, current deep-learning-facilitated FLFM methods encounter fundamental limitations. Training typically relies on externally acquired high-resolution ground-truth volumes, which impose an experimental burden and limit scalability. Moreover, their performance is often tightly coupled to the specific sample or imaging conditions employed for training, necessitating labor-intensive retraining or fine-tuning when applied to varying specimen types, morphologies, or dynamic regimes. These limitations underscore the need for a broadly generalizable and accessible framework for 3D light-field reconstructions.

In this work, we introduce PAVR, a physics-aware, universal Fourier light-field volumetric reconstruction framework for high-resolution cellular imaging. PAVR minimizes reliance on experimental training data and on external high-end modalities by training exclusively on *in-silico* point-spread functions (PSFs). Without imposing prior assumptions about sample conditions, PAVR learns the intrinsic optical response of the FLFM system, enabling robust, generalizable spatiotemporal reconstruction across a wide range of biological contexts. We demonstrate the versatility of PAVR through multicolor volumetric imaging of subcellular structures in fixed and live cells, three-dimensional autofluorescence particle tracking, and quantitative analysis of morphological and functional dynamics in beating human cardio-myocytes under pharmacological perturbation. Together, these results establish PAVR as a scalable and widely applicable framework that accelerates high-throughput volumetric imaging across basic cell biology and translational applications.

## RESULTS

### PAVR architecture and implementation

Unlike existing frameworks, PAVR features physics-guided supervision with numerically generated datasets and high-speed volumetric reconstruction with high generalizability. The PAVR framework is trained exclusively on *in silico*, physics-aware datasets that explicitly encode a wave-optics forward model of FLFM. This design eliminates dependence on external experimental training data and obviates sample-specific retraining by exploiting the unified, system-defined PSF intrinsic to FLFM (**Supplementary Table 1**). Generalizability across diverse biological structures relies on the fact that all fluorescence images can be represented as superpositions of sample-independent point emitters, each governed by the system PSF (**Fig. 1a and Methods**). Building on this principle, we implement a cascaded training data synthesis strategy that spans a broad range of spatial organizations and emitter densities, including uniform distribution, Neyman-Scott aggregation^51^, clustering, and random-walk-based continuous distributions^52^. These synthesized datasets comprehensively recapitulate experimentally observed fluorescence patterns while mitigating sample dependence, thereby enabling robust reconstruction across heterogeneous 3D features (**Methods**). In addition, PAVR incorporates a physically informed supervision scheme (**Fig. 1b, c**) that explicitly mitigates defocus-induced blur in the synthesized training data. This constraint suppresses artifacts caused by axial elongation and supports accurate 3D reconstruction of structurally complex cellular morphologies (**Methods**). The resulting paired wide-field (WF) stacks and light-field (LF) images derived from unified PSFs serve as physics-consistent ground truth (GT) and network inputs, allowing the model to learn a stable and generalizable inverse mapping from LF measurements to volumetric reconstructions (**Fig. 1d and Supplementary Fig. 1**).

**Figure 1.**
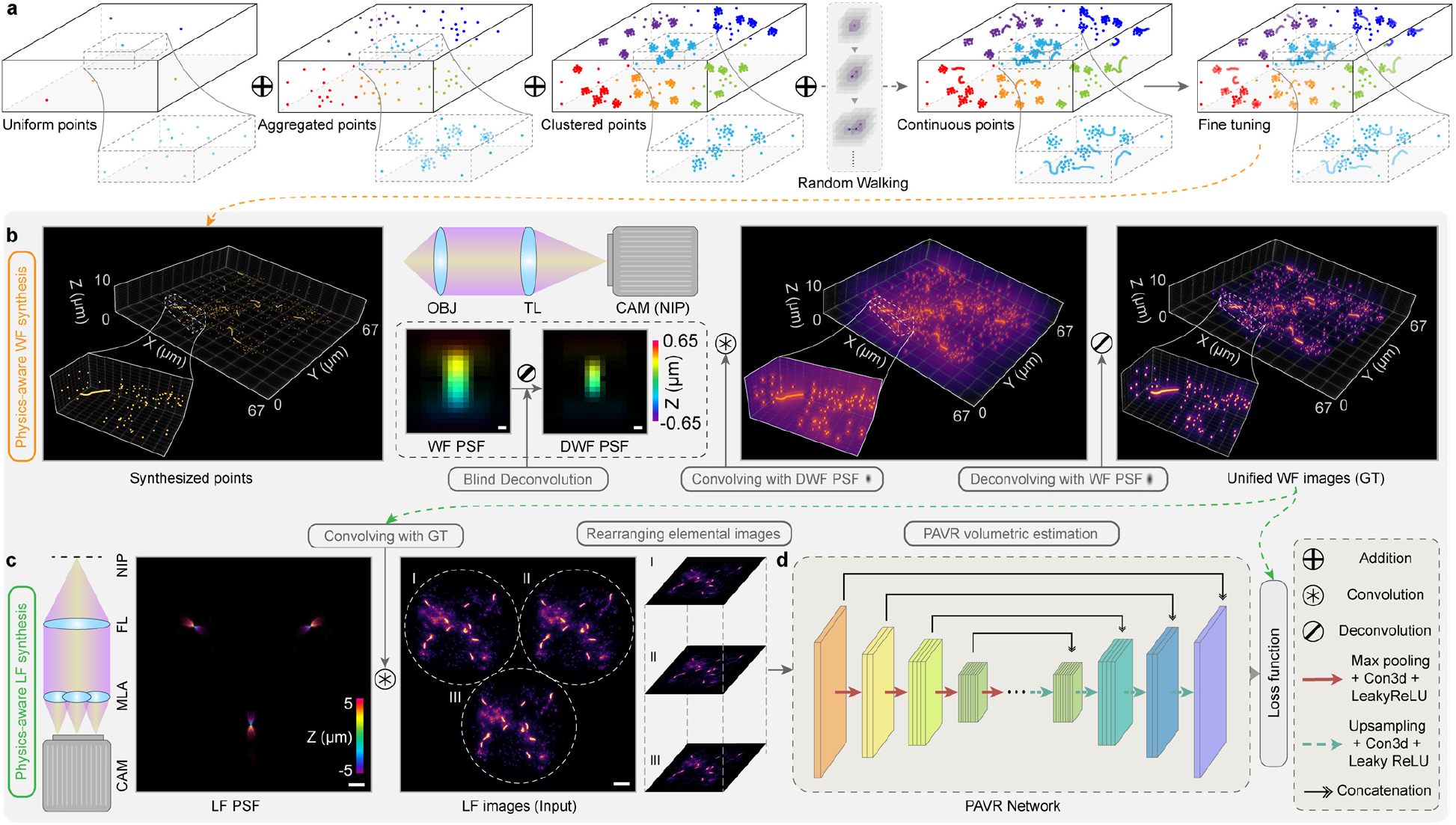
Physics-Aware Volumetric Reconstruction Framework (PAVR). **(a)** Physics-aware synthesis of three-dimensional distributed, sub-diffraction–limited microspheres from randomly initialized single-pixel emitters. Uniformly distributed points are first generated throughout the volume to define a coarse scaffold. Aggregated emitters are then formed around each point by sampling from anisotropic Gaussian distributions, followed by deriving clustered emitters to introduce finer-scale micro-heterogeneity. Continuous emitter distributions are generated by linking points to produce spatially connected trajectories. Intensities are assigned and normalized via connectivity-based analysis to ensure consistent brightness across structures of varying size and density, and a mild Gaussian blur with intensity adjustment yields realistic, continuous, and intensity-balanced 3D bead distributions for training and validation. **(b)** Synthesis of wide-field (WF) ground-truth volumes. The simulated bead distributions in (a) are convolved with a blind-deconvolved WF point-spread function (PSF) and subsequently deconvolved using the original WF PSF to generate high-fidelity volumetric ground truth. **(c)** Generation of Fourier light-field (LF) network inputs. The WF ground-truth volumes in (b) are convolved with a Fourier light-field PSF to form LF images, which are then segmented into elemental images and axially rearranged to construct the network input tensor. **(d)** Training workflow. Paired WF ground truth and LF inputs are used to train a 3D U-Net–based model, optimized using mean squared error loss. Scale bars: 10 µm.

### High-resolution imaging of subcellular organelles in mammalian cells

To experimentally validate the performance of the PAVR framework, we imaged multiple subcellular organelles across distinct cell types with diverse structural complexity. We first captured Fourier light-field images of GFP-labeled peroxisomes in HeLa cells (**Fig. 2a-c, Methods**). The elemental images were processed with PAVR, which exhibited near-diffraction-limited resolution of approximately 350 nm and 600 nm in the lateral and axial directions, respectively. The results substantially suppressed reconstruction artifacts and preserved fine 3D structural details, compared with conventional Richardson-Lucy deconvolution^53^ and wide-field images (**Fig. 2d, e**, and **Supplementary Fig. 2**). To assess network generalizability across cellular contexts and morphologies, we next imaged immunostained microtubules in U2OS cells (**Fig. 2f, g, Supplementary Fig. 3, and Methods**). Quantitative cross-sectional analyses revealed consistent diffraction-limited resolutions (**Fig. 2h-j**). In addition, PAVR results showed markedly enhanced contrast and metrics compared to conventional deconvolution and wide-field images (**Fig. 2k**). Furthermore, we demonstrated the framework using two-color imaging of mitochondria and nuclei in HeLa cells (**Fig. 2l-o and Supplementary Fig. 4**). Unlike peroxisomes or microtubules, these organelles exhibit extended, heterogeneous morphologies and broad axial distributions, placing greater demands on volumetric reconstruction. Despite this complexity, PAVR maintained high spatial resolution and reconstruction fidelity, with minimal artifacts and higher SNR than deconvolution (**Fig. 2p**). Analysis of mitochondrial and nuclear images verified consistent diffraction-limited resolution in all three dimensions and robust 3D two-color image registration with enhanced clarity^54,55^ (**Fig. 2q, r, Supplementary Table 2, and Supplementary Note 1**).

**Figure 2.**
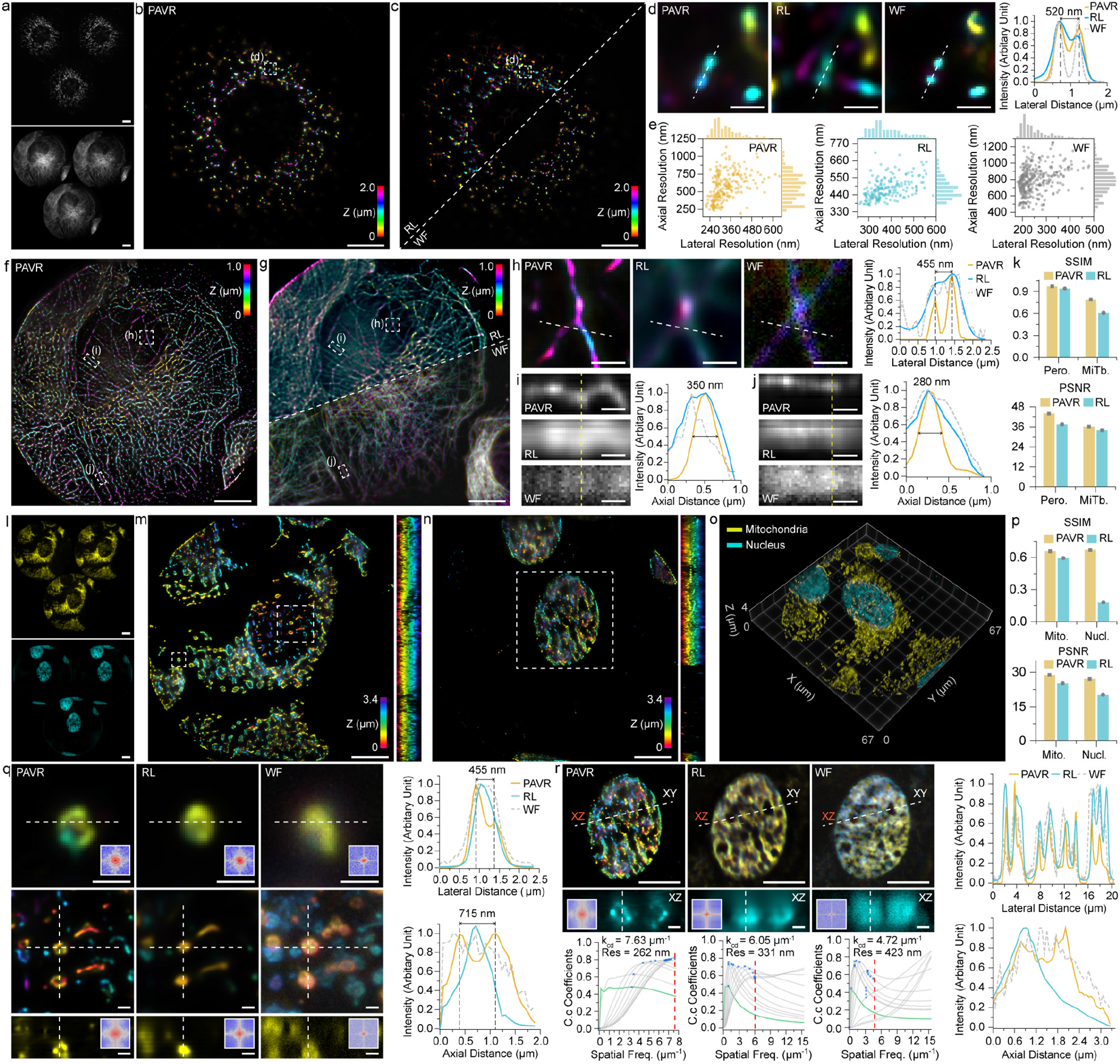
High-resolution PAVR imaging of subcellular organelles in mammalian cells. **(a)** Denoised Fourier light-field (LF) images of GFP-labeled peroxisomes in HeLa cells (top) and immunostained microtubules in U2OS cells (bottom). **(b, c)** Corresponding 3D reconstructions of peroxisomes (a, top) using PAVR (b), in comparison with Richardson–Lucy deconvolution (RL; c, top left), and wide-field axial scanning (WF; c, bottom right). **(d)** Zoomed-in images of the boxed regions in (b, c) with cross-sectional profiles, showing the resolution of nearby structures separated by 520 nm with PAVR. **(e)** Distributions of 3D Gaussian-fitted peroxisome FWHMs for PAVR (yellow, N = 220), RL (cyan, N = 214), and WF (gray, N = 341). PAVR preserves a high diffraction-limited resolution consistent with RL and WF. **(f, g)** Corresponding 3D reconstructions of microtubules (a, bottom) using PAVR (f), in comparison with RL (g, top), and WF (g, bottom). **(h)** Zoomed-in images of the boxed regions in (f, g) with cross-sectional profiles, showing the resolution of nearby filaments separated by 455 nm with PAVR. **(i, j)** Corresponding axial views of the boxed regions in (f, g) with cross-sectional profiles (dashed lines), demonstrating enhanced axial FWHM measurements of individual filaments at approximately 300 nm. **(k)** Improved reconstruction fidelity of PAVR images over RL, in quantitative comparison of SSIM and PSNR versus WF images. **(l)** Denoised two-color LF images of mitochondria (top) and nuclei (bottom) in HeLa cells. **(m, n)** Corresponding color-coded 3D PAVR reconstructions of mitochondria (m) and nuclei (n). **(o)** 3D view of the merged two-color reconstructions in (m, n). **(p)** Comparison of SSIM and PSNR for mitochondria and nuclei, indicating enhanced reconstruction with PAVR. **(q)** Zoomed-in images of the boxed regions in (m) comparing PAVR with RL and WF. Fourier-spectrum analysis (insets) in lateral and axial dimensions, along with corresponding profiles, indicate ring structures separated by 455 nm laterally and 715 nm axially with PAVR. **(r)** Zoomed-in images of the boxed regions in (n) comparing PAVR with RL and WF. Fourier-spectrum analysis (insets) and decorrelation-based resolution estimation (bottom) yield resolutions of 262 nm (PAVR), 331 nm (RL), and 423 nm (WF), with corresponding 3D cross-sectional profiles shown at right. Scale bars: 10 µm (a-c, f, g, l-n, r top), 1 µm (d, h-j, q, r bottom).

### Live-cell imaging with high-spatiotemporal resolution

We next performed live-cell imaging and assessed PAVR for dynamic volumetric reconstruction. We initially imaged autofluorescent lysosomes in cardiac fibroblasts derived from human-induced pluripotent stem cells (hiPSCs) at an acquisition of 100 ms per volume using 488-nm excitation (**Fig. 3a** and **Supplementary Movie 1**). For lysosome tracking, PAVR exhibited isotropic localization, providing approximately 300 nm resolution for 3D lysosomal trajectories, and rapid volumetric rendering enabled near-instant 3D volume visualization (approximately 10 Hz volume rate) during acquisition (**Fig. 3b, c**). This high spatiotemporal resolution facilitated multiparticle transport and interaction analysis, allowing direct observation of lysosomal fission and fusion events, processes central to organelle homeostasis, cargo processing, and metabolic and signaling regulation^56,57^ (**Fig. 3d**). Furthermore, we demonstrated PAVR under reduced signal-to-noise conditions and higher temporal demands. We conducted two-color live-cell imaging of mitochondria and the Golgi apparatus in bone marrow-derived mesenchymal stem cells (MSCs), whose interactions are crucial for studying energy supply, protein trafficking, and many organelle-related diseases, such as Alzheimer’s^58^. Employing stroboscopic illumination at 200 Hz, we acquired dual-color volumetric images in each frame at an effective frame rate of 100 Hz over a continuous 20-s acquisition period^41^ (**Fig. 3e-p, Supplementary Fig. 5, Supplementary Movie 2**, and **Methods**). Across an axial range of 4 µm, PAVR consistently recovered both organelles with improved resolution and axial fidelity compared to conventional deconvolution (**Fig. 3f-i**). Notably, the high volumetricity and spatiotemporal resolution enabled by PAVR allowed for clear visualization of rapid mitochondrial morphological remodeling in three dimensions and volumetric interactions between mitochondria and the Golgi apparatus (**Fig. 3j-l and Supplementary Fig. 6**).

**Figure 3.**
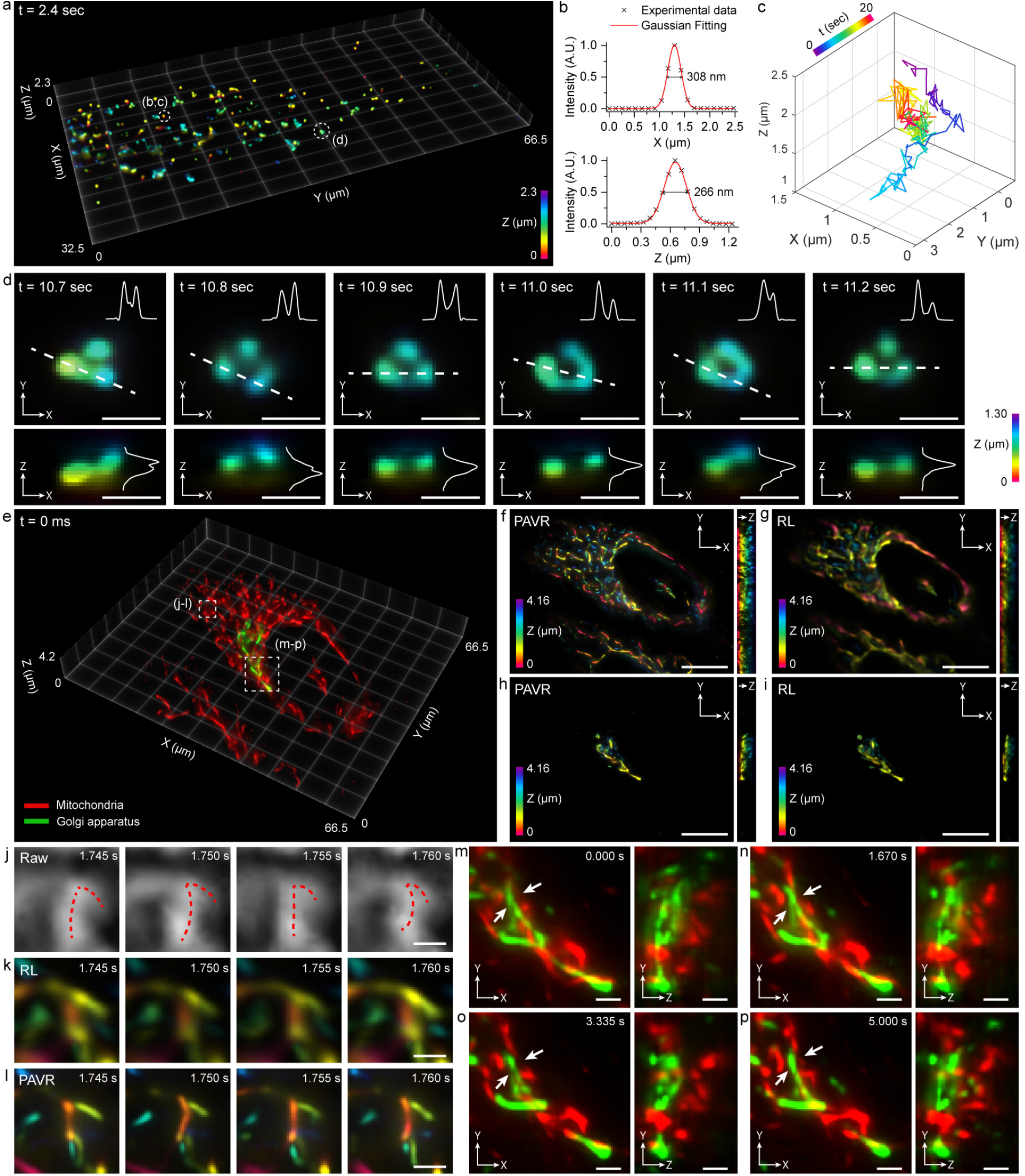
Live-cell PAVR imaging with high-spatiotemporal resolution. **(a)** 3D visualization of lysosomal autofluorescence in live hiPSC-derived fibroblasts acquired at 10 volumes per second. **(b)** Representative localization profiles of a single lysosomal particle (a) fitted with a 3D Gaussian model, exhibiting 3D FWHM values of approximately 300 nm. **(c)** 3D trajectory of the lysosomal particle tracked over 20 s at 10 Hz. **(d)** 3D views of time-resolved morphological remodeling of lysosomal clusters (a) at 10 Hz. **(e)** 3D two-color rendering of mitochondria (red) and Golgi apparatus (green) in live bone marrow–derived mesenchymal stem cells (MSCs) acquired using stroboscopic illumination for a volume rate at 200 Hz (effective 100 Hz per color). **(f-i)** Color-coded 3D maximum-intensity projections of mitochondria (f,g) and Golgi (h,i) reconstructed with PAVR (f,h) and Richardson–Lucy (RL) deconvolution (g,i). **(j-l)** Zoomed-in raw LF (j), corresponding RL (k), and PAVR (l) images of mitochondria in the boxed region in (e) at four time points. **(m-p)** Zoomed-in 3D views of the two-color reconstructions at four time points, displaying rapid volumetric remodeling and organelle interactions. Scale bars: 10 µm (f-i), 1 µm (d, j-p).

### Time-lapse imaging of pharmacologically induced morphological dynamics in cardiomyocytes

Lastly, we demonstrated sensitive, high-speed volumetric detection of cardiomyocyte (CM) morphological responses to pharmacological perturbation (**Fig. 4, Supplementary Fig. 7**, and **Supplementary Movies 3, 4**). Isoproterenol (ISO), a β-adrenergic agonist clinically used to treat cardiac conduction disorders such as bradycardia and heart block, increases cardiomyocyte (CM) beating frequency, contractility, and vasodilatation^59,60^ through mechanisms that critically depend on mitochondrial function and cellular energy homeostasis^61,62^. At elevated doses, however, ISO induces oxidative stress, leading to alterations in mitochondrial membrane potential (MMP) and potentially causing cardiac injury^63^. Because both mitochondrial morphology and MMP evolve dynamically within the subcellular environment, accurate characterization of ISO-induced responses requires high-resolution, high-speed volumetric imaging with minimal reconstruction artifacts.

**Figure 4.**
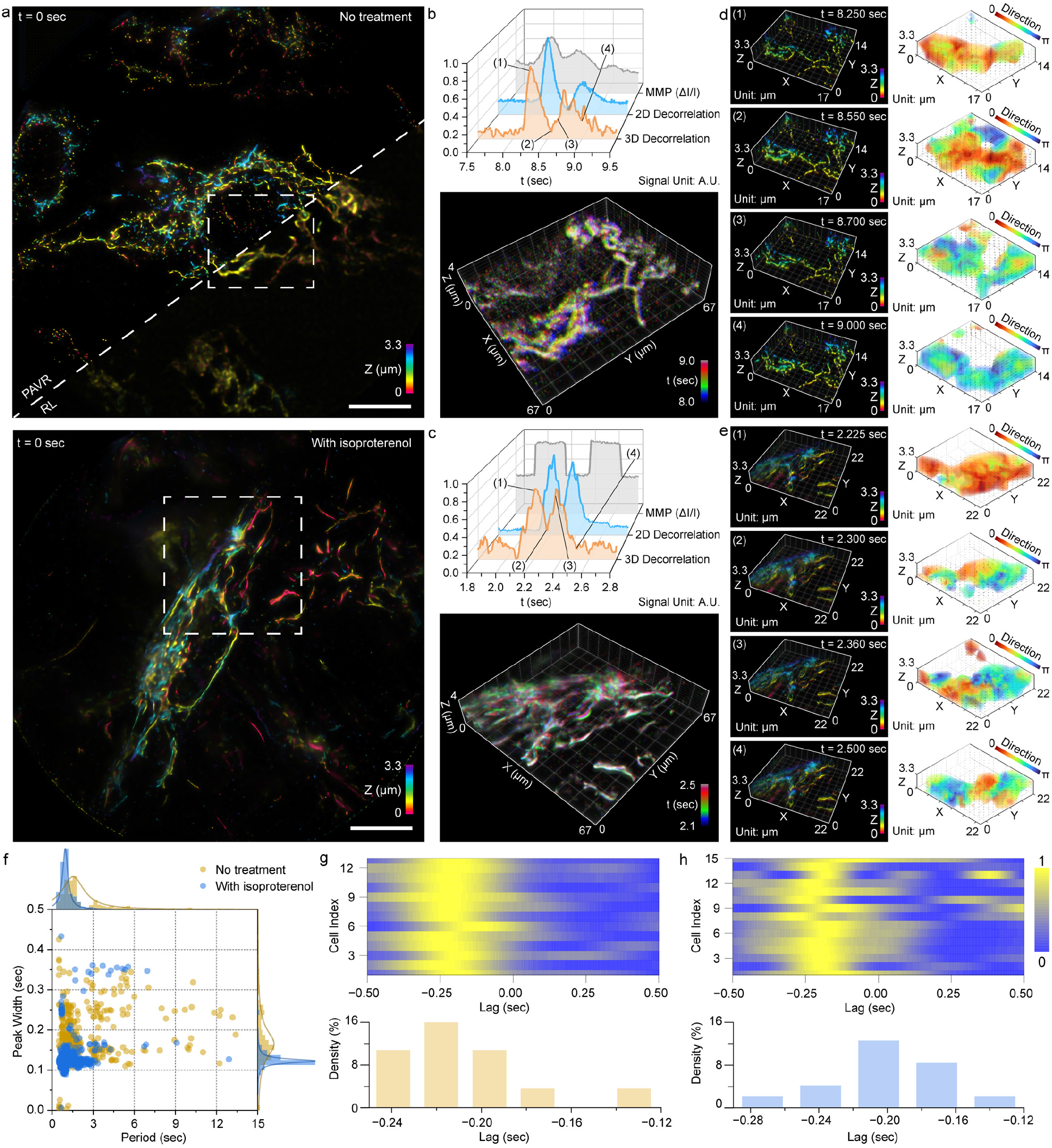
Time-lapse PAVR imaging of pharmacologically induced morphological dynamics in cardiomyocytes. **(a)** Depth-coded images of mitochondria in hiPSC-derived cardiomyocytes (CMs) reconstructed with PAVR under untreated (top) and isoproterenol (ISO)–treated (bottom) conditions. Exemplary RL reconstruction is shown for comparison. **(b, c)** Top panels, representative beating traces extracted from mitochondrial membrane-potential (TMRM intensity) signals and from morphology-derived spatial-correlation signals for untreated (b) and ISO-treated (c) CMs. Bottom panels, montage of time-coded 3D volumes of the boxed regions in (a), exhibiting time-resolved mitochondrial dynamics during beating in untreated (b) and ISO-treated (c) CMs. **(d, e)** Left panels, PAVR reconstructions of the boxed untreated (d) and ISO-treated (e) mitochondrial volumes at four time points. Right panels, corresponding 3D displacement vector fields computed from the reconstructions, with colors indicating displacement direction relative to the initial contraction direction. **(f)** Distributions of beating period and peak width extracted from morphology-derived traces for untreated (yellow, N = 409 beats) and ISO-treated (blue, N = 1161 beats) CMs, revealing a more intensive CM beating with ISO treatment. **(g, h)** Cross-correlation analysis between morphology-derived and membrane-potential–derived traces as a function of temporal lag (top; normalized) for untreated (g, N = 13 cells) and ISO-treated (h, N = 15 cells) CMs, with corresponding lag histograms (bottom), showing an average delay of ∼200 ms of morphology-derived signals relative to membrane-potential–derived traces. Scale bars: 10 µm.

To this end, we imaged untreated and ISO-treated hiPSC-derived CMs, with mitochondria labeled with the MMP-sensitive dye tetramethyl rhodamine methyl ester (TMRM) at an acquisition rate of 200 Hz over 1 min (**Methods**). PAVR captured both mitochondrial morphology and intensity-based MMP dynamics. Across conditions, PAVR captured both mitochondrial morphology and intensity-based MMP dynamics, and the results consistently outperformed conventional deconvolution in terms of spatial resolution and noise suppression (**Fig. 4a and Supplementary Fig. 7**). Temporal analysis revealed that morphology-derived signals robustly captured CM beating dynamics with substantially higher signal-to-noise ratio than MMP-derived intensity traces, despite both modalities reporting the same underlying contractile events (**Fig. 4b, c, and Supplementary Fig. 8**). This indicates that volumetric morphological readouts provide a more precise and reliable basis for beat detection and quantification^64^. Compared with 2D projections, the 3D volumes produced by PAVR yielded pixel-wise displacement maps across all three dimensions, directly revealing subcellular patterns of mitochondrial networks during CM contraction cycles (**Fig. 4d, e**). In particular, the analysis of 13 untreated and 15 ISO-treated hiPSC-CMs exhibited that ISO treatment significantly increased CM beating frequency while producing faster and more abbreviated contraction profiles, consistent with prior reports^65^ (**Fig. 4f**). Notably, joint volumetric analysis of MMP and morphology further revealed that morphological deformation followed MMP changes with a delay of 211 ± 30 ms for untreated cells and 193 ± 43 ms for ISO-treated cells, an observation in agreement with previous physiological measurements using minimum-quadratic-difference-based particle image velocimetry^66^ (**Fig. 4g, h**). These results highlight PAVR as a broadly applicable volumetric reconstruction framework that bridges subcellular structural architecture with quantitative functional dynamics, which opens new opportunities for cardiac disease modeling^67^, drug-induced cardiotoxicity assessment^68^, and patient-specific functional phenotyping for precision pharmacology^69^.

## DISCUSSION

Modern biological microscopy is increasingly shaped by a hardware-software co-design paradigm, in which optical encoding and computational reconstruction are jointly required to deliver high-resolution, volumetric imaging at biologically relevant speeds. Emerging FLFM techniques provide an efficient optical route to single-shot volumetric acquisition; however, their broad utility in cell biology has been constrained by the computational burden, limited throughput, and reconstruction artifacts associated with iterative model-based approaches. Here, we address these challenges by introducing PAVR, a physics-aware volumetric reconstruction framework that fuses hardware-facilitated volumetric acquisition with fast, artifact-suppressed computational inference, thereby enabling practical high-resolution 3D imaging across diverse cellular contexts. PAVR overcomes a central limitation of learning-assisted light-field reconstruction by employing a sample-independent training strategy that does not rely on external high-resolution modalities for ground truth (**Supplementary Note 2**). The current implementation adopts an accessible 3D U-Net architecture and achieves an approximately 100-fold acceleration over conventional iterative deconvolution while maintaining superior reconstruction quality (**Supplementary Note 3, Supplementary Movie 5**). Moreover, the PVAR framework is readily extensible to alternative U-Net variants, including residual^70^ and attention-augmented^71^ architectures, and is compatible with emerging hardware configurations^72,73^ (**Supplementary Note 4**). The enhanced capabilities of PAVR will enable high-throughput and population-scale applications, including imaging flow cytometry^41,74^, image-based cell profiling^75,76^, and single-molecule analysis^77^ (**Methods, Supplementary Note 5)**. We anticipate that PAVR will serve as a versatile paradigm for advancing high-resolution, high-throughput volumetric imaging across fundamental cell biology and translational research domains.

## METHODS

### Fourier light-field microscopy

The PAVR imaging experiments were performed on a previously developed high-resolution FLFM using an epi-fluorescence microscope (Eclipse Ti2-U, Nikon Instruments)^53^. In the microscope system, we implemented a 100× oil-immersion objective lens with a numerical aperture of 1.45 (CFI Plan Apochromat Lambda 100× Oil, Nikon Instruments) and a piezo nano-positioner (Nano-F100S, Mad City Labs) for precise positioning. During the experiments, samples were excited using multicolor laser lines (405 nm, Coherent; 488 nm, 561 nm, 647 nm, MPB Communications), while the fluorescence was collected through a quadband dichroic mirror (ZT405/488/561/647, Chroma) and a corresponding emission filter (ZET405/488/561/647 m, Chroma). A micro-positioning system (MS2000, Applied Scientific Instrumentation) was used for accurate sample positioning. A Fourier lens Fourier-transformed the native image plane of the objective lens to the image pupil (*f*_FL_ = 275 mm, Edmund Optics). A customized microlens array (*f*_ML_ = 117 mm, RPC Photonics) was positioned on the image pupil to form the elemental images, which were acquired by an sCMOS camera (ORCA-Flash 4.0 V3, Hamamatsu Photonics, pixel size *P*_cam_ = 6.5 µm).

### Light-field image acquisition

The fixed-cell imaging experiments were performed with a camera exposure time of 100 ms. We first captured a high-resolution Fourier light-field image of the sample, then switched the microscope’s observation portal to the other side for wide-field scanning without moving the samples. The wide-field scanning was performed by shifting the piezo positioner in an axial range of 10 µm with a step size of 65 nm. For live-cell imaging, we put the sample in a stage-top incubator with an objective heater, digital gas mixing, and digital sensor-driven humidity (H301, Okolab). We used a camera exposure time of 100 ms for fibroblast autofluorescence imaging and 5 ms for all other live-cell imaging experiments. In particular, we implemented a stroboscopic illumination scheme for two-color live-cell imaging, alternating between the two laser lines in each camera exposure period^41^. Therefore, each fluorescence was captured at a camera exposure of 5 ms and a frame rate of 100 Hz.

### Light-field image preprocessing

During training, we adjusted all clear WF and LF images to random intensity values within the signal power range specified by the training parameters. In the predictions, the experimental light-field images were first denoised using ACsN^78^ and MIRO^79^ algorithms. For images of complex, thick samples, an additional rolling-ball background-subtraction approach^80^ was implemented to remove defocus blurs. Empirically, for 16-bit greyscale images with a maximum intensity of at least 1000, the robustness of network testing and prediction is demonstrated without any denoising schemes.

### Generating wave-optics-based training datasets

The PSF generation process began with the generation of initial points, which were uniformly distributed across the 3D volume (1024 × 1024 × 155 voxels, voxel size of 65 nm), to define the global spatial distribution of the VPs. These points served as the coarse structural basis, establishing large-scale organization and depth variation within the simulated field. To further improve network performance and make it more adaptive to diverse structures, we intentionally add Neyman-Scott PSF clusters with varying densities^51^ and random-walk-based, continuously positioned PSFs^52^ in addition to the initial virtual points (**Fig. 1a** and **Methods**). Specifically, the aggregated points were generated around each initial point by sampling from an anisotropic Gaussian distribution with varying densities and extents that mimic realistic bead aggregation patterns. A further refinement step produced clustered points, derived from a subset of the aggregated points to form finer-scale micro-clusters and introduce multi-level spatial heterogeneity. To enhance connectivity, continuous points were generated via single-pixel biased random walks that favor lateral expansion, creating filament-like trajectories and continuous spatial transitions. After the spatial structure was established, each point was assigned to random intensity values, followed by normalization through density analysis to ensure consistent brightness across clusters and strings of varying sizes. Finally, the entire volume was applied a mild Gaussian blur to round bead edges, resulting in a realistic and intensity-balanced 3D sub-diffraction-limited point distribution suitable for training and validation in light-field imaging simulations.

The wide-field (WF) and the corresponding light-field (LF) images are then synthesized by convolving with a WF PSF (1024 × 1024 × 155 voxels with a voxel size of 65 nm) and an LF PSF (1024 × 1024 × 155 voxels with a voxel size of 145 nm), respectively. In the defocus compensation scheme, we first refine the original WF PSF *h*_0_ using a blind deconvolution procedure to effectively yield a sharpened PSF *h*_s_, which partially inverts the residual defocus and optical aberrations. In the frequency domain, this corresponds to *H*_s_ = *H*_0_/*G*(ω), where *H*_s_ and *H*_0_ are the Fourier transforms of *h*_s_ and *h*_0_, respectively, *G*(ω) represents the defocus-induced low-pass modulation transfer function, and ω represents spatial frequency. Next, the clear WF images are generated by a forward convolution with *h*_s_ and a subsequent Richardson-Lucy deconvolution with *h*_0_ (**Fig. 1b**), while the corresponding clear LF images are generated by a forward convolution between a hybrid LF PSF (hPSF)^53^ and the clear WF images (**Fig. 1c**). Therefore, the WF transfer function *H*_WF_ equals 1/*G*(ω) and the equivalent LF transfer function *H*_eq_ equals *H*_LF_/*G*(ω), where *H*_LF_ is the Fourier transform of the hPSF. Hence, the *H*_eq_ effectively compensates for defocus blur and improves focus fidelity. Moreover, the training dataset was augmented through lateral translations and axial rotations. These transformations helped create a robust training set by simulating various sample orientations and positions within the field of view. In practice, we generated 80 pairs of training images and augmented each pair 4 times, yielding a total of 400 pairs.

### PAVR network architecture

The architecture of PAVR was specifically designed for reconstructing Fourier LF images (**Supplementary Fig. 1**). The network input received Fourier LF images, with each elemental image rearranged into different channels of the tensor. In the network, the input images were first expanded in the axial direction using a 2D convolutional layer to align with the desired network output size. An additional dimension was then inserted to reposition the previous channel dimension to the axial dimension, which was compatible with the 3D U-Net scenario. Subsequently, the 3D tensors were processed by an initial channel-repetition layer for further feature extraction in the 3D U-Net architecture. The 3D U-Net architecture contained 7 layers with both encoding and decoding pathways. The encoding steps (red solid arrows in **Fig. 1d**) consisted of 3D convolutional blocks that progressively extracted the spatial dimensions of the feature maps into the channel dimension. Each encoding layer implemented 3D convolutions followed by LeakyReLU non-linearities, capturing increasingly abstract features of the input LF images. The high-dimensional features were compressed into a lower-dimensional spatial representation at the network’s bottleneck, capturing the features necessary for accurate 3D reconstructions. The decoding steps (green dashed arrows in **Fig. 1d**) reversed the encoding pathway, consisting of up-sampling layers that increased the spatial dimensions of the feature maps, followed by convolutional layers with non-linear LeakyReLU activation functions that refined the features. This pathway reconstructed the 3D spatial information from the encoded volumes. Skip connections (black double arrows in **Fig. 1d** and **Supplementary Fig. 1**) linked corresponding layers in the encoding and decoding pathways, allowing the network to retain high-resolution features and prevent the loss of spatial information. These connections facilitated the direct flow of information and gradients, improving the network’s ability to learn and reconstruct fine details. The final convolutional layer, followed by a squeezing layer, produced the high-resolution 3D reconstructed image for subsequent analysis and visualization of single cells.

### PAVR network training

Before sending the ground truth (GT) and network input (NI) pairs into the network, the NI images were segmented, resized, padded, and rearranged into different channels. Specifically, a 1024-by-1024-pixel light-field image with a 145-nm pixel size was segmented into three 500-by-500 elemental images with a 145-nm pixel size. The elemental images were then resized and padded to 1024-1024 pixels at a pixel size of 65 nm, consistent with the GT voxel size, using bicubic interpolation and zero-padding. The transformed elemental images were then rearranged into a 1024-by-1024-by-3-voxel stack with a voxel size of 65 nm. Limited by GPU memory (48 GB, Nvidia A6000), both GT and NI datasets were laterally rescaled by a factor of 0.5, and the GT depth was trimmed to 64 layers. The final GT volume size was 512 × 512 × 64 voxels, and the NI image stack size was 512 × 512 × 3 voxels. During the training process, the mean squared error (MSE) and structural similarity index (SSIM)^81^ were used to evaluate the accuracy of the 3D reconstruction. Empirically, the learning rate was exponentially reduced from the 160th epoch to reduce overfitting. A dropout rate of 0.3 and batch normalization were also used to further improve the network’s generalizability and stability during training. Typically, the network can be well-trained with 300-400 epochs.

### Culturing HeLa and U2OS cell lines

HeLa and U2OS cells were cultured according to the same protocol. In short, cells were cultured in DMEM with 10% FBS and 1% Penicillin-streptomycin (Pen-Strep) under 37 °C and 5% CO_2_ conditions (**Supplementary Table 3**). Once the cell cultures reached ∼80% confluency, cells were passaged and cultured into culture dishes or imaging dishes, depending on the needs.

### Labeling peroxisomes in live HeLa cells

Peroxisome labeling was performed with live HeLa cells. On the day prior to the imaging session, cells were passaged to a 35 mm imaging dish and cultured in DMEM with 20 µL of CellLight peroxisome-GFP solution (**Supplementary Table 3**). Cells were then incubated at 37 °C and 5% CO_2_ conditions for 18 hours. On the imaging day, cells were washed 2 rounds with DMEM, and the dish was immediately used for imaging.

### Immunostaining of microtubules in fixed U2-OS cells

Fixed microtubule staining was performed with U2-OS cells. Cells were passaged and cultured in Lab-Tek II chambered slides containing DMEM in each well until the desired confluency was reached. For immunostaining, each well on the slide was first washed once with PBS. Then, each well was fixed with 3% PFA and 0.1% glutaraldehyde in PBS at room temperature for 12 minutes. Extra aldehyde groups were reduced with 0.1% sodium borohydride for 7 minutes, followed by 3 rounds of PBS washing for 5 minutes each. After that, cells were permeabilized and blocked with a blocking buffer (3% BSA with 0.2% Triton X-100 in PBS) for 30 minutes at room temperature. Cells were incubated for 30 minutes with β-tubulin primary antibody (BT7R, 6 µg/mL) diluted in blocking buffer at room temperature. Next, each well was washed 5 times with washing buffer (0.2% BSA with 0.05% Triton X-100 in PBS) for 15 minutes per wash at room temperature. After washing, labeled secondary antibody dilutions (goat anti-mouse, Alexa Fluor 647, 3 µg/mL) in blocking buffer were added to each well and incubated for 30 minutes at room temperature, with light avoided (**Supplementary Table 3**). Then, each well was washed 3 times with the washing buffer for 10 minutes per wash at room temperature, followed by a 5-minute wash in PBS. Lastly, cells were fixed again with 3% PFA and 0.1% glutaraldehyde at room temperature for 10 minutes, followed by 3 rounds of PBS washes for 5 minutes each. Cells were stored in PBS for imaging purposes.

### Two-color labeling of the nucleus and mitochondria in fixed HeLa cells

Two-color staining was performed with fixed HeLa cells. HeLa cells were passaged and cultured in a chambered slide with DMEM in each well until the desired confluency was reached. The mitochondria immunostaining was performed similarly to the above microtubule immunostaining protocol; however, the primary antibody was switched to TOMM20 (3G5, 4 µg/mL), and the secondary antibody was switched to an anti-rabbit Alexa Fluor 647-conjugated antibody (3 µg/mL). The conditions for the remaining steps of the immunostaining protocol (i.e., fixation, washing, blocking, permeabilization, and antibody incubation) remained the same. After the second round of cell fixation, the slide was stained with DAPI dilutions (1 µg/mL) for 5 minutes (**Supplementary Table 3**). Each well was rinsed with PBS, and the final chambered slide was stored in PBS.

### Preparation of cardiac fibroblasts induced from human-induced pluripotent stem cells (hiPSCs)

MR90-IV (WiCell Research Institute) was maintained in mTeSR1 medium (Stem Cell Technologies) and induced for cardiac differentiation as previously described^82^. Briefly, hiPSCs were treated with 100 µg/mL recombinant human activin A (R&D Systems) in insulin-free RPMI medium containing 2% B27 (RPMI-B27) on day 0. After 24 h, the medium was replaced with 10 µg/mL recombinant human bone morphogenic protein-4 (BMP4, R&D Systems) in RPMI-B27 insulin-free medium from day 1 to day 4. The medium was changed to RPMI medium with 2% B27 containing insulin (RPMI-B27 medium) on day 4. Cardiac spheroids were generated on day 10 post-differentiation. Differentiated cells were dissociated with 0.25% trypsin–EDTA and seeded into AggreWell 400 plates (number 34415, Stem Cell Technologies) at 1,500 cells per microwell to form cardiac spheroids. Before cell seeding, plates containing 1 ml of RPMI–B27 medium per well were centrifuged at 1000 g to remove trapped bubbles in the microwells. To prevent cell death, the medium was supplemented with 10 μM of Rock inhibitor Y-27632 (Selleck Chemicals). Plates were centrifuged at 100 g to distribute the cells and then placed in an incubator. After 24 h, spheroids were collected to remove the Rock inhibitor, and the RPMI-B27 medium was replaced in the suspension culture. Three-dimensional spheroids were maintained in poly-D-lysine (FD35PDL-100, FluoroDish) to promote cardiac fibroblast proliferation at the dish surface. Cultures were maintained until the day of testing (<30 days post-differentiation). The medium was refreshed every 4-5 days.

### Culturing and staining of bone marrow-derived mesenchymal stem cells (MSCs)

MSCs (#PCS-500-012, ATCC) were maintained in Dulbecco’s Modified Eagle Medium (DMEM; #10-013-CV, Corning) supplemented with 10% fetal bovine serum (FBS; #35-011-CV, Corning) and 1% Penicillin–Streptomycin (#15140122, ThermoFisher). Cells were cultured at 37 °C in a humidified incubator with 5% CO_2_. Before imaging, the Golgi apparatus was stained with BODIPY Ceramide–BSA Complexes (#B22650, ThermoFisher), and the mitochondria were stained with MitoTracker Deep Red (M22426, ThermoFisher). The 5-µM BODIPY staining solution was prepared by diluting the stock solution 100-fold (e.g., adding 10 µL of stock solution to 1 mL of PBS). The MitoTracker staining solution was prepared by diluting 1-mM MitoTracker stock solution to the final working concentration of 25-500 nM. The cells were first rinsed twice with PBS on a 37 °C heating pad and incubated for 30 min at 4°C with 5 µM ceramide-BSA complexes. After staining the Golgi apparatus, the sample was rinsed again with cold medium and incubated in fresh DMEM at 37 °C. At this step, we added 2 mL of MitoTracker working solution and incubated for 30 min. After the staining was complete, the cells were rinsed twice with PBS and stored in fresh prewarmed media for imaging experiments.

### Preparation, staining, and imaging of hiPSC-CMs

The hiPSCs (IMR90, WiCell Research Institute) were maintained on Matrigel-coated 12-well plates in mTeSR1 medium (Stem Cell Technologies) with daily media changes until reaching ∼70-80% confluence. After 3-4 days of daily medium changes, when cultures reached 90-100% confluence, cells were treated with RPMI 1640 supplemented with 2% B27 minus insulin (B27(-)) and 1% penicillin-streptomycin (RPMI+B27(-)+P/S) containing 6 µM CHIR99021 (S1263, SelleckChem) for 48 h, followed by 24 h of RPMI+B27(-)+P/S without CHIR to initiate cardiomyocyte differentiation. Subsequently, 5 µM IWR-1 (I0161-5MG, Sigma-Aldrich) was applied in RPMI+B27(-)+P/S for 48 h. Cultures were then maintained in RPMI supplemented with B27 plus insulin (B27(+)), and P/S (RPMI+B27(+)+P/S), with media refreshed every 2-3 days until cardiomyocyte differentiation was evident.

On the day of imaging, mitochondrial membrane potential was assessed using tetramethylrhodamine methyl ester (TMRM, T668, ThermoFisher). A staining solution was prepared by diluting 2 µL of 100 µM TMRM stock into 2 mL of prewarmed culture medium, yielding a final concentration of 100 nM. Then, the original maturation medium of the cells was replaced with the staining medium, and the cells were incubated in the incubator for 45 min. After that, the staining medium was replaced with 2 mL prewarmed maturation medium for recovery, and the cells were incubated for 30 min or more. Videos lasting 60 s were taken at 200 Hz for each hiPSC-CM sample.

### Chemical and biological materials

The sources of the chemicals and biological materials used in the experiments, including company names and catalog numbers, are listed in **Supplementary Table 3**. No further authentication procedures are conducted after receiving the stock cell lines directly from the suppliers.

### Statistics and reproducibility

The fluorescence staining protocol was repeated at least three times for each experiment. During data acquisition, samples were imaged at least 5 times per experiment. Images with optimal fluorescence brightness were selected for the figures.

## DATA AVAILABILITY

Example data and network model checkpoints are available at https://doi.org/10.5281/zenodo.18446762. The data underlying the results presented in this paper can be obtained from the corresponding author upon request due to the large file size. Requests will be fulfilled within two weeks. Source data is provided with this paper.

## CODE AVAILABILITY

The code package for the PAVR is available as Supplementary Software. The code has been written in MATLAB 2025a (MathWorks) and Python 3.11. The latest version of the software will be made available upon publication at: https://github.com/ShuJiaLab/PAVR.

## ACKNOWLEDGEMENTS

We acknowledge support from National Science Foundation grants 2503686, 2225990, and 2145235, National Institutes of Health grants R35GM124846 and R21HD110918, and the faculty start-up fund of the Georgia Institute of Technology.

## AUTHOR CONTRIBUTIONS STATEMENT

X.H. and S.J. conceived and designed the project. X.H. and K.H. contributed to the construction of the imaging system. X.H. developed deep-learning algorithms. K.H. and W.L. helped with HeLa cells. K.H. helped with U2OS cells. Z.L., O.R., P.F., and C.X. helped with hiPSC-induced fibroblasts and cardio-myocytes. Z.G., H.Z., and E.A.B. helped with bone marrow-derived mesenchymal stem cells. M.A.S. and A.K. helped with H460 cell samples. A.R., H.K., and J.E.D. helped with mouse endothelial cells. X.H. and K.H. prepared and labeled samples. X.H. performed imaging experiments and analyzed the data. J.E.D., A.K., E.A.B., and C.X. contributed scientific insights. S.J. supervised the overall project. X.H. and S.J. wrote the manuscript with input from all authors.

## REFERENCES

1 Balasubramanian, H., Hobson, C. M., Chew, T. L. & Aaron, J. S. Imagining the future of optical microscopy: everything, everywhere, all at once. Commun Biol 6, 1096 (2023). 10.1038/s42003-023-05468-9

2 Hockenberry, M. A., Daugird, T. A. & Legant, W. R. Cell dynamics revealed by microscopy advances. Curr Opin Cell Biol 90, 102418 (2024). 10.1016/j.ceb.2024.102418

3 Morris, J. D. & Payne, C. K. Microscopy and Cell Biology: New Methods and New Questions. Annu Rev Phys Chem 70, 199–218 (2019). 10.1146/annurev-physchem-042018-052527

4 Wang, Z., Peng, Y., Fang, L. & Gao, L. Computational optical imaging: on the convergence of physical and digital layers. Optica 12, 113–130 (2025). 10.1364/OPTICA.544943

5 Shroff, H., Testa, I., Jug, F. & Manley, S. Live-cell imaging powered by computation. Nature Reviews Molecular Cell Biology 25, 443–463 (2024). 10.1038/s41580-024-00702-6

6 Belthangady, C. & Royer, L. A. Applications, promises, and pitfalls of deep learning for fluorescence image reconstruction. Nat Methods 16, 1215–1225 (2019). 10.1038/s41592-019-0458-z

7 Mengu, D. et al. At the intersection of optics and deep learning: statistical inference, computing, and inverse design. Advances in Optics and Photonics 14, 209–290 (2022). 10.1364/Aop.450345

8 Archana, R. & Jeevaraj, P. S. E. Deep learning models for digital image processing: a review. Artificial Intelligence Review 57, 11 (2024). 10.1007/s10462-023-10631-z

9 Cheng, H. C. et al. Deep-Learning-Assisted Volume Visualization. IEEE Trans Vis Comput Graph 25, 1378–1391 (2019). 10.1109/TVCG.2018.2796085

10 Moen, E. et al. Deep learning for cellular image analysis. Nat Methods 16, 1233–1246 (2019). 10.1038/s41592-019-0403-1

11 Lopez-Jimenez, A. T. et al. High-content high-resolution microscopy and deep learning-assisted analysis reveals host and bacterial heterogeneity during Shigella infection. Elife 13 (2025). 10.7554/eLife.97495

12 Ma, Q. & Xu, D. Deep learning shapes single-cell data analysis. Nat Rev Mol Cell Biol 23, 303–304 (2022). 10.1038/s41580-022-00466-x

13 Chai, B., Efstathiou, C., Yue, H. & Draviam, V. M. Opportunities and challenges for deep learning in cell dynamics research. Trends Cell Biol 34, 955–967 (2024). 10.1016/j.tcb.2023.10.010

14 Zhang, Y. Z. & Imoto, S. Genome analysis through image processing with deep learning models. J Hum Genet 69, 519–525 (2024). 10.1038/s10038-024-01275-0

15 Kobayashi, H., Cheveralls, K. C., Leonetti, M. D. & Royer, L. A. Self-supervised deep learning encodes high-resolution features of protein subcellular localization. Nat Methods 19, 995–1003 (2022). 10.1038/s41592-022-01541-z

16 Pratapa, A., Doron, M. & Caicedo, J. C. Image-based cell phenotyping with deep learning. Curr Opin Chem Biol 65, 9–17 (2021). 10.1016/j.cbpa.2021.04.001

17 Sapoval, N. et al. Current progress and open challenges for applying deep learning across the biosciences. Nat Commun 13, 1728 (2022). 10.1038/s41467-022-29268-7

18 Serag, A. et al. Translational AI and Deep Learning in Diagnostic Pathology. Front Med (Lausanne) 6, 185 (2019). 10.3389/fmed.2019.00185

19 Rivenson, Y. et al. Deep Learning Enhanced Mobile-Phone Microscopy. ACS Photonics 5, 2354–2364 (2018). 10.1021/acsphotonics.8b00146

20 Park, B. et al. Rapid cancer diagnosis using deep learning-powered label-free subcellular-resolution photoacoustic histology. Sci Adv 11, eadz1820 (2025). 10.1126/sciadv.adz1820

21 Wang, S. et al. Towards next-generation diagnostic pathology: AI-empowered label-free multiphoton microscopy. Light Sci Appl 13, 254 (2024). 10.1038/s41377-024-01597-w

22 Nehme, E., Weiss, L. E., Michaeli, T. & Shechtman, Y. Deep-STORM: super-resolution single-molecule microscopy by deep learning. Optica (2018). 10.1364/optica.5.000458

23 Zhang, M. et al. Deep learning enhanced light sheet fluorescence microscopy for in vivo 4D imaging of zebrafish heart beating. Light Sci Appl 14, 92 (2025). 10.1038/s41377-024-01710-z

24 Qiao, C. et al. Rationalized deep learning super-resolution microscopy for sustained live imaging of rapid subcellular processes. Nat Biotechnol 41, 367–377 (2023). 10.1038/s41587-022-01471-3

25 Guan, H. et al. Deep-learning two-photon fiberscopy for video-rate brain imaging in freely-behaving mice. Nat Commun 13, 1534 (2022). 10.1038/s41467-022-29236-1

26 Guo, M. et al. Deep learning-based aberration compensation improves contrast and resolution in fluorescence microscopy. Nat Commun 16, 313 (2025). 10.1038/s41467-024-55267-x

27 Wang, Z. et al. Real-time volumetric reconstruction of biological dynamics with light-field microscopy and deep learning. Nature Methods 18 (2021). 10.1038/s41592-021-01058-x

28 Sheneman, L., Balogun, S., Johnson, J. L., Harrison, M. J. & Vasdekis, A. E. Near-zero photon bioimaging by fusing deep learning and ultralow-light microscopy. Proc Natl Acad Sci U S A 122, e2412261122 (2025). 10.1073/pnas.2412261122

29 Zhao, R. et al. A review of light-field imaging in biomedical sciences. Med X 3, 25 (2025). 10.1007/s44258-025-00070-6

30 Llavador, A., Sola-Pikabea, J., Saavedra, G., Javidi, B. & Martínez-Corral, M. Resolution improvements in integral microscopy with Fourier plane recording. Optics Express 24, 20792–20792 (2016). 10.1364/oe.24.020792

31 Scrofani, G. et al. FIMic: design for ultimate 3D-integral microscopy of in-vivo biological samples. Biomedical Optics Express 9, 335–335 (2018). 10.1364/BOE.9.000335

32 Cong, L. et al. Rapid whole brain imaging of neural activity in freely behaving larval zebrafish (Danio rerio). eLife 6, e28158–e28158 (2017). 10.7554/eLife.28158

33 Guo, C., Liu, W., Hua, X., Li, H. & Jia, S. Fourier light-field microscopy. Optics Express 27, 25573–25594 (2019). 10.1364/OE.27.025573

34 Sims, R. R. et al. Single molecule light field microscopy. Optica 7, 1065–1065 (2020). 10.1364/OPTICA.397172

35 Daly, S. et al. High-density volumetric super-resolution microscopy. Nat Commun 15, 1940 (2024). 10.1038/s41467-024-45828-5

36 Li, H. et al. Fast, volumetric live-cell imaging using high-resolution light-field microscopy. Biomedical Optics Express 10, 29–29 (2019). 10.1364/boe.10.000029

37 Hua, X., Liu, W. & Jia, S. High-resolution Fourier light-field microscopy for volumetric multi-color live-cell imaging. Optica (2021). 10.1364/optica.419236

38 Han, K. et al. 3D super-resolution live-cell imaging with radial symmetry and Fourier light-field microscopy. Biomed. Opt. Express 13, 5574–5584 (2022). 10.1364/BOE.471967

39 Ling, Z., Han, K., Liu, W., Hua, X. & Jia, S. Volumetric live-cell autofluorescence imaging using Fourier light-field microscopy. Biomedical Optics Express 14, 4237–4245 (2023).

40 Wu, J. et al. Iterative tomography with digital adaptive optics permits hour-long intravital observation of 3D subcellular dynamics at millisecond scale. Cell 184, 3318–3332 e3317 (2021). 10.1016/j.cell.2021.04.029

41 Hua, X. et al. Light-field flow cytometry for high-resolution, volumetric and multiparametric 3D single-cell analysis. Nat Commun 15, 1975 (2024). 10.1038/s41467-024-46250-7

42 Liu, W., Kim, G. R., Takayama, S. & Jia, S. Fourier light-field imaging of human organoids with a hybrid point-spread function. Biosens Bioelectron 208, 114201 (2022). 10.1016/j.bios.2022.114201

43 Ling, Z. et al. Multiscale and recursive unmixing of spatiotemporal rhythms for live-cell and intravital cardiac microscopy. Nat Cardiovasc Res 4, 637–648 (2025). 10.1038/s44161-025-00649-7

44 Zhou, K. C. et al. High-speed 4D fluorescence light field tomography of whole freely moving organisms. Optica 12, 674–684 (2025). 10.1364/optica.549707

45 Levoy, M., Zhang, Z. & McDowall, I. Recording and controlling the 4D light field in a microscope using microlens arrays. Journal of Microscopy 235, 144–162 (2009). 10.1111/j.1365-2818.2009.03195.x

46 Fu, B. et al. Patch deconvolution for Fourier light-field microscopy. bioRxiv, 2025.2006.2013.659385 (2025).

47 Lin, B., Tian, Y., Zhang, Y., Zhu, Z. & Wang, D. Deep learning methods for high-resolution microscale light field image reconstruction: a survey. Front Bioeng Biotechnol 12, 1500270 (2024). 10.3389/fbioe.2024.1500270

48 Vizcaino, J. P. et al. Real-time light field 3D microscopy via sparsity-driven learned deconvolution. 2021 IEEE International Conference on Computational Photography (ICCP), 1–11 (2021).

49 Yi, C. Q. et al. Video-rate 3D imaging of living cells using Fourier view-channel-depth light field microscopy. Communications Biology 6 (2023). 10.1038/s42003-023-05636-x

50 Zhu, L. et al. Adaptive-learning physics-assisted light-field microscopy enables day-long and millisecond-scale super-resolution imaging of 3D subcellular dynamics. Nat Commun 16, 7132 (2025). 10.1038/s41467-025-62471-w

51 Wang, Y., Degleris, A., Williams, A. & Linderman, S. W. Spatiotemporal Clustering with Neyman-Scott Processes via Connections to Bayesian Nonparametric Mixture Models. J Am Stat Assoc 119, 2382–2395 (2024). 10.1080/01621459.2023.2257896

52 Altendorf, H. & Jeulin, D. Random-walk-based stochastic modeling of three-dimensional fiber systems. Phys Rev E Stat Nonlin Soft Matter Phys 83, 041804 (2011). 10.1103/PhysRevE.83.041804

53 Hua, X., Liu, W. & Jia, S. High-resolution Fourier light-field microscopy for volumetric multi-color live-cell imaging. Optica 8, 614–620 (2021). 10.1364/optica.419236

54 Descloux, A., Grußmayer, K. S. & Radenovic, A. Parameter-free image resolution estimation based on decorrelation analysis. Nature Methods 16, 918–924 (2019). 10.1038/s41592-019-0515-7

55 Cang, H., Liu, Y. & Xing, J. Mosaic-PICASSO: accurate crosstalk removal for multiplex fluorescence imaging. Bioinformatics 40 (2024). 10.1093/bioinformatics/btad784

56 Ballabio, A. & Bonifacino, J. S. Lysosomes as dynamic regulators of cell and organismal homeostasis. Nat Rev Mol Cell Biol 21, 101–118 (2020). 10.1038/s41580-019-0185-4

57 Li, X. et al. A molecular mechanism to regulate lysosome motility for lysosome positioning and tubulation. Nat Cell Biol 18, 404–417 (2016). 10.1038/ncb3324

58 Crawford, K. et al. Golgi apparatus, endoplasmic reticulum and mitochondrial function implicated in Alzheimer’s disease through polygenic risk and RNA sequencing. Mol Psychiatry 28, 1327–1336 (2023). 10.1038/s41380-022-01926-8

59 Zou, Y. et al. Isoproterenol Activates Extracellular Signal–Regulated Protein Kinases in Cardiomyocytes Through Calcineurin. Circulation 104, 102–108 (2001). doi:10.1161/hc2601.090987

60 Denis, A. et al. Diagnostic Value of Isoproterenol Testing in Arrhythmogenic Right Ventricular Cardiomyopathy. Circulation: Arrhythmia and Electrophysiology 7, 590–597 (2014). doi:10.1161/CIRCEP.113.001224

61 Dedkova, E. N. & Blatter, L. A. Measuring mitochondrial function in intact cardiac myocytes. J Mol Cell Cardiol 52, 48–61 (2012). 10.1016/j.yjmcc.2011.08.030

62 Liu, L. et al. The interplay between cardiac dyads and mitochondria regulated the calcium handling in cardiomyocytes. Front Physiol 13, 1013817 (2022). 10.3389/fphys.2022.1013817

63 Mukherjee, D. et al. Mechanisms of isoproterenol-induced cardiac mitochondrial damage: protective actions of melatonin. J Pineal Res 58, 275–290 (2015). 10.1111/jpi.12213

64 Pardon, G. et al. Tracking single hiPSC-derived cardiomyocyte contractile function using CONTRAX an efficient pipeline for traction force measurement. Nat Commun 15, 5427 (2024). 10.1038/s41467-024-49755-3

65 Hansen, K. J. et al. Optical Method to Quantify Mechanical Contraction and Calcium Transients of Human Pluripotent Stem Cell-Derived Cardiomyocytes. Tissue Eng Part C Methods 23, 445–454 (2017). 10.1089/ten.TEC.2017.0190

66 Ahola, A., Polonen, R. P., Aalto-Setala, K. & Hyttinen, J. Simultaneous Measurement of Contraction and Calcium Transients in Stem Cell Derived Cardiomyocytes. Ann Biomed Eng 46, 148–158 (2018). 10.1007/s10439-017-1933-2

67 Lewis-Israeli, Y. R. et al. Self-assembling human heart organoids for the modeling of cardiac development and congenital heart disease. Nat Commun 12, 5142 (2021). 10.1038/s41467-021-25329-5

68 Schwach, V. et al. A safety screening platform for individualized cardiotoxicity assessment. iScience 27, 109139 (2024). 10.1016/j.isci.2024.109139

69 Aikawa, M. et al. Precision cardiovascular medicine: shifting the innovation paradigm. Front Sci 3 (2025). 10.3389/fsci.2025.1474469

70 He, K., Zhang, X., Ren, S. & Sun, J. in Proceedings of the IEEE conference on computer vision and pattern recognition. 770–778 (2016).

71 Oktay, O. et al. Attention U-Net: Learning Where to Look for the Pancreas. arXiv preprint arXiv:1804.03999 (2018). 10.48550/arXiv.1804.03999

72 Hsieh, H.-C. et al. Imaging 3D cell cultures with optical microscopy. Nature Methods 22, 1167–1190 (2025). 10.1038/s41592-025-02647-w

73 Zhao, R. et al. A review of light-field imaging in biomedical sciences. Med-x 3, 25 (2025).

74 Ling, Z. et al. LFC-plus: simultaneous multicolour volume cytometry for high-throughput single-cell analysis. Lab on a Chip (2026).

75 Caicedo, J. C. et al. Data-analysis strategies for image-based cell profiling. Nature Methods 14, 849–863 (2017). 10.1038/nmeth.4397

76 Nitta, N. et al. Clinical-grade autonomous cytopathology through whole-slide edge tomography. Nature (2026). 10.1038/s41586-025-10094-y

77 Han, K. et al. Volumetric localization microscopy with deep learning. Nat Commun 16, 10960 (2025). 10.1038/s41467-025-65941-3

78 Mandracchia, B. et al. Fast and accurate sCMOS noise correction for fluorescence microscopy. Nat Commun 11, 94 (2020). 10.1038/s41467-019-13841-8

79 Mandracchia, B. et al. Optimal sparsity allows reliable system-aware restoration of fluorescence microscopy images. Sci Adv 9, eadg9245 (2023). 10.1126/sciadv.adg9245

80 Sternberg, S. R. Biomedical Image-Processing. Computer 16, 22–34 (1983).

81 Wang, Z., Bovik, A. C., Sheikh, H. R. & Simoncelli, E. P. Image quality assessment: from error visibility to structural similarity. IEEE transactions on image processing 13, 600–612 (2004).

82 Ling, Z., Han, K., Liu, W., Hua, X. & Jia, S. Volumetric live-cell autofluorescence imaging using Fourier light-field microscopy. Biomed Opt Express 14, 4237–4245 (2023). 10.1364/BOE.495506

